# Phage-Encoded LuxR-Type Receptors Responsive to Host-Produced Bacterial Quorum-Sensing Autoinducers

**DOI:** 10.1101/577726

**Authors:** Justin E. Silpe, Bonnie L. Bassler

## Abstract

Quorum sensing (QS) is a process of cell-to-cell communication that bacteria use to orchestrate collective behaviors. QS relies on the cell-density-dependent production, accumulation, and receptor-mediated detection of extracellular signaling molecules called autoinducers (AIs). Gram-negative bacteria commonly use *N*-acyl homoserine lactones (AHLs) as their AIs and they are detected by LuxR-type receptors. Often, LuxR-type receptors are insoluble when not bound to a cognate AI. In this report, we show that LuxR-type receptors are encoded on phage genomes and, in the cases we tested, the phage LuxR-type receptors bind to and are solubilized specifically by the AHL AI produced by the host bacterium. We do not yet know the viral activities that are controlled by these phage QS receptors, however, our observations, coupled with recent reports, suggest that their occurrence is more widespread than previously appreciated. Using receptor-mediated detection of QS AIs could enable phages to garner information concerning the population density status of their bacterial hosts. We speculate that such information can be exploited by phages to optimize the timing of execution of particular steps in viral infection.

**Importance:** Bacteria communicate with chemical signal molecules to regulate group behaviors in a process called quorum sensing (QS). In this report, we find that genes encoding receptors for Gram-negative bacterial QS communication molecules are present on genomes of viruses that infect these bacteria. These viruses are called phages. We show that two phage-encoded receptors, like their bacterial counterparts, bind to the communication molecule produced by the host bacterium, suggesting that phages can “listen in” on their bacterial hosts. Interfering with bacterial QS and using phages to kill pathogenic bacteria represent attractive possibilities for development of new antimicrobials to combat pathogens that are resistant to traditional antibiotics. Our findings of interactions between phages and QS bacteria need consideration as new antimicrobial therapies are developed.

---

Bacteria engage in the cell-cell communication process called quorum sensing (QS) to coordinate group behaviors. QS relies on production, release, and group-wide detection of extracellular signal molecules called autoinducers (AIs). QS-mediated-communication systems are now known to be common in the bacterial domain (1). Growing evidence suggests that viruses of bacteria, called phages, also possess communication capabilities, an idea that originated when Hargreaves et al. reported the presence of genes resembling those encoding Gram-positive bacterial QS components (*agr*) on a phage infecting *Clostridium difficile* (2). More recent studies show that phages can engage in their own QS-like dialogs to promote population-wide lysogeny (3), and phages can eavesdrop on bacterial QS-mediated communication to promote population-wide lysis (4). In the eavesdropping case, which is most pertinent to the observations we report here, a vibriophage encodes a homolog of a *Vibrio* QS receptor/transcription factor called VqmA. Host-encoded VqmA and phage-encoded VqmA (VqmA_Phage_) both bind the same AI, 3,5-dimethylpyrazin-2-ol (DPO). In the case of the host, binding of DPO by VqmA launches the QS group behavior program (5). In the case of the phage, binding of DPO by VqmA_Phage_ activates the lysis program and the phage kills the host. Thus, the vibriophage lysis-lysogeny decision is linked to the population-density status of the host cells (4).

Inspired by the above findings, we wondered whether additional phage-bacterial QS connections could exist. To explore this possibility, we performed database analyses scanning for QS receptors encoded by viral genomes. We focused on genes encoding *N*-acyl homoserine lactone (AHL)-binding LuxR-type receptors/transcription factors because they exist in thousands of sequenced genomes of Gram-negative bacteria, making them an easily recognizable type of QS AI-receptor pair. Our bioinformatic search for receptors belonging to the “transcription factor LuxR-like, AI-binding domain” superfamily of proteins (InterPro: IPR036693) revealed 12,285 AHL-binding LuxR domain entries encoded by bacteria and two additional entries in a category outside of living organisms, among viruses. One hit was present on a putative *Myoviridiae* phage (accession: MH622937.1) in metagenomic data from animal viruses (Christopher Buck NIH; personal communication; Figure S1A). Because this DNA sequence is not a verified phage and the phage does not exist in a known or available host bacterial strain, we could not pursue this finding further. The second viral *luxR*-type gene was on a characterized phage that infects *Aeromonas sp*. ARM81 (accession: KT898133.1). This phage is called ΦARM81ld (6) (Figure 1A) and the gene designated *p37* encodes a putative LuxR-type transcription factor. While we were unable to obtain *Aeromonas sp*. ARM81 or the resident phage, we synthesized the *p37* gene and cloned it into a recombinant expression vector. Hereafter, we call the protein encoded by *p37* LuxR_ΦARM81ld_.

**Figure 1.**
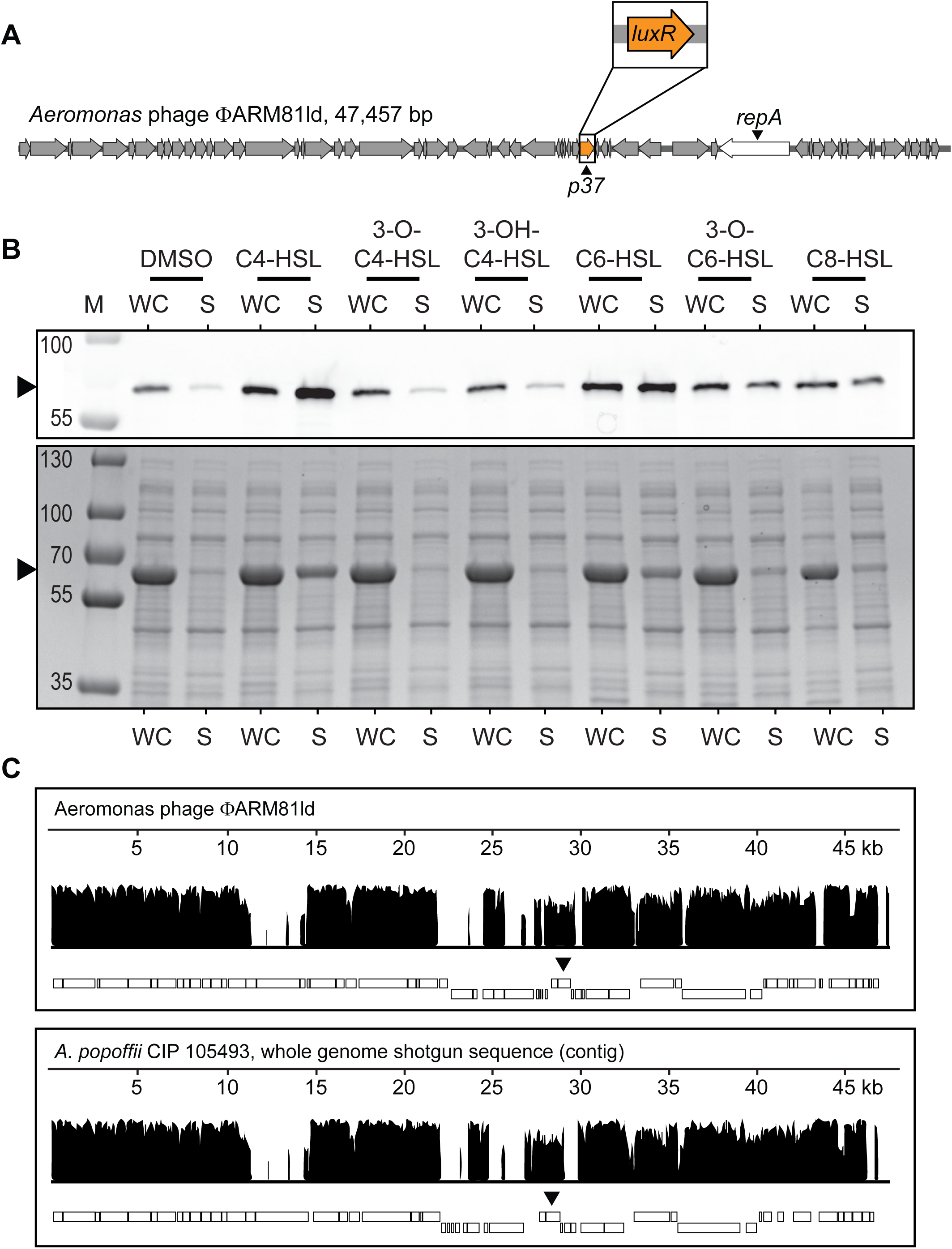
Phage ΦARM81ld encodes a QS LuxR-type receptor that is solubilized by bacterial AIs and *A. popoffii* is lysogenized by a phage-like element similar to ΦARM81ld. (A) Genome organization of phage ΦARM81ld. *p37* is predicted to encode a Lux-type receptor (orange). The phage replication gene *repA* is displayed in white. (B) Western blot (upper) and total protein (lower) showing HALO-LuxR_ΦARM81ld_ in the whole cell (WC) lysate and soluble (S) fractions of recombinant *E. coli* supplied with 75 µM of the indicated AHL or an equivalent volume of DMSO. Arrowheads indicate HALO-LuxR_ΦARM81ld_. M denotes marker (PageRuler Plus; representative bands are labeled). Regarding the differences in band intensities, Coomassie stains total protein, folded and unfolded, whereas the HALO-Western technique requires that the HALO-tag be functional for detection. Thus, the HALO-LuxR_ΦARM81ld_ bands in the WC samples show only the fraction of the protein that is folded and functional, and thus they appear fainter than in the corresponding bands in the S fractions that show all of the folded and functional protein. Consistent with this interpretation, the Coomassie stained gel shows that there is indeed more total HALO-LuxR_ΦARM81ld_ present in the WC than the S fractions. (C) Whole phage-genome alignment of the known phage ΦARM81ld against the full-length *A. popoffii* phage-like contig. Nucleotide sequence identity (vertical axis) along the length of the aligned genomes (horizontal axis). The rectangles below each plot represent predicted ORFs. Upper and lower boxes represent the + and – strands, respectively, on which the ORFs are encoded. The black arrows indicate the locations of the phage *luxR* genes. Alignment performed using the progressive Mauve algorithm (see Methods).

Commonly, bacterial LuxR-type receptors require their cognate AHL ligands to fold and, in turn, become soluble (7). To determine whether the phage-encoded LuxR_ΦARM81ld_ receptor follows the same constraints, we expressed HALO-tagged LuxR_ΦARM81ld_ in *E. coli*. SDS-PAGE and Western Blot with HALO-Alexa_660_ showed that the LuxR_ΦARM81ld_ protein was produced and present in the whole cell lysate but not in the soluble fraction (Figure 1B). We reasoned that LuxR_ΦARM81ld_ might bind the C4-homoserine lactone (C4-HSL) AI produced by members of the host *Aeromonas* genus (8). Indeed, Figure 1B shows that addition of C4-HSL to the *E. coli* carrying HALO-LuxR_ΦARM81ld_ enabled the HALO-LuxR_ΦARM81ld_ to fold and become soluble as it was present in both the whole cell lysate and in the soluble fraction. To examine ligand specificity, we assessed the ability of a panel of AHL AIs with varying tail lengths (C4, C6, C8) and decorations (3-OH, 3-O) to solubilize LuxR_ΦARM81ld_. LuxR_ΦARM81ld_ is solubilized by C4-HSL but not by AHLs with C4 tails containing decorations (Figure 1B). AHLs containing acyl tails longer than C4 or a longer tail and a decoration (3-O-C6-HSL) display diminished abilities to solubilize LuxR_ΦARM81ld_ compared to C4-HSL (Figure 1B). These results demonstrate that, like bacterial LuxR-type receptors which often show exquisite specificity for their cognate AHL AIs, the phage-encoded LuxR_ΦARM81ld_ receptor is solubilized primarily by the AHL AI reportedly produced by its *Aeromonad* host genus.

There are no homologs identical to LuxR_ΦARM81ld_ in the NCBI database, however, LuxR_ΦARM81ld_ shares ∼48% identity with the bacterial QS receptor encoded by the human pathogen *Aeromonas hydrophila* called AhyR (8) and ∼60% homology to a LuxR-type receptor on a DNA contig from an uncharacterized isolate of *Aeromonas popoffii* CIP 105493 (9). Curiously, *A. popoffii* CIP 105493 has at least two genes encoding putative LuxR receptors (Figure S1B). The first *A. popoffii* homolog, with 48% identity to LuxR_ΦARM81ld_, shares 95% identity with *A. hydrophila* AhyR and is located in the same genomic context as *ahyR* in *A. hydrophila* (neighboring a putative *luxI* gene called *ahyI*), indicating that this first homolog is likely the *A. popoffii* bacterial QS receptor. The second *A. popoffii* homolog has less than 50% identity to *A. hydrophila* AhyR but, as mentioned, shares ∼60% identity to LuxR_ΦARM81ld_ (Figure S1B). This gene is located on a ∼47 kb contig that, by PHASTER analysis (10), is predicted to be an intact phage, suggesting that like LuxR_ΦARM81ld_, this second *A. popoffii luxR* gene exists on a phage. Consistent with this idea, we found that, as is the case for *Aeromonas sp*. ARM81 (6), lysis of *A. popoffii* is induced by the DNA damaging agent mitomycin C (MMC). Specifically, a precipitous decline in OD_600_ over time occurs following MMC addition (Figure S2A). DNA specific to the phage-like contig can be amplified from the clarified supernatants of DNase-treated culture lysates, suggesting that phage particles are present. Furthermore, PCR amplification of the putative phage origin and primase gene required for replication (*repA*), ligation of the product to an antibiotic resistance cassette followed by introduction and selection in *E. coli*, shows that the ligated product can be maintained as a plasmid. This final result demonstrates that the *luxR*-containing contig encodes a functional replication gene for a plasmid-like element, which is the reported state in which ΦARM81ld exists when it lysogenizes its *Aeromonas* host (6). Lastly, the contig carrying this putative phage *luxR* gene is comparable in length to the genome of phage ΦARM81ld (46.8 kb in *A. popoffii* vs. 47.6 kb in ΦARM81ld) and can be aligned along its entire length to the complete ΦARM81Id genome (60.1% pairwise identify; Figure 1C). Thus, we hypothesize that *A. popoffii* harbors an extrachromosomally-replicating prophage encoding a *luxR* QS gene similar to ΦARM81ld. We hereafter refer to the phage in *A. popoffii* as Apop and the phage LuxR receptor as LuxR_Apop_. Similar to the vibriophage-*Vibrio* case described above (4), because of the low sequence identity, the QS receptors encoded on ΦARM81ld and on Apop do not appear to be the result of direct transfer from the bacterial host.

The predicted AhyR-like QS receptors encoded by the *A. popoffii* and *Aeromonas sp*. ARM81 bacterial hosts share more than 90% identity, and yet the corresponding phage-encoded receptors LuxR_Apop_ and LuxR_ΦARM81ld_, respectively, are only ∼60% identical to one another (Figures 2A, S2B, and S2C). This disparity in amino acid identity between the phage LuxR proteins led us to wonder whether the LuxR_Apop_ AI binding preferences were the same or different from that of LuxR_ΦARM81ld_. Of the seven AHLs in our test collection, recombinant LuxR_Apop_ is solubilized by C4-HSL and also by C6- and C8-HSL, but it is not solubilized by AHLs containing tail decorations (Figure 2B). Consistent with the idea that the two phage receptors recognize the AI produced by their bacterial hosts, cell-free culture fluids from *A. popoffii* activated an *E. coli* reporter strain that generates light specifically in response to exogenously supplied C4-HSL (Figure 2C) and LC-MS analysis of the *A. popoffii* culture fluid showed that C4-HSL was present (Figure 2D) in amounts comparable to those reported for other C4-HSL-producing bacteria (>10 µM by both assays) (11). We could not quantify C6- or C8-HSL in the *A. popoffii* fluids, suggesting that those molecules are either not made by *A. popoffii* or they exist below the limit of our method (∼500 and 250 nM for C6- and C8-HSL, respectively). Taken together, our results suggest that, like ΦARM81ld, the Apop phage encodes a functional LuxR receptor that binds to and is solubilized by C4-HSL, an AI produced by its bacterial host.

**Figure 2.**
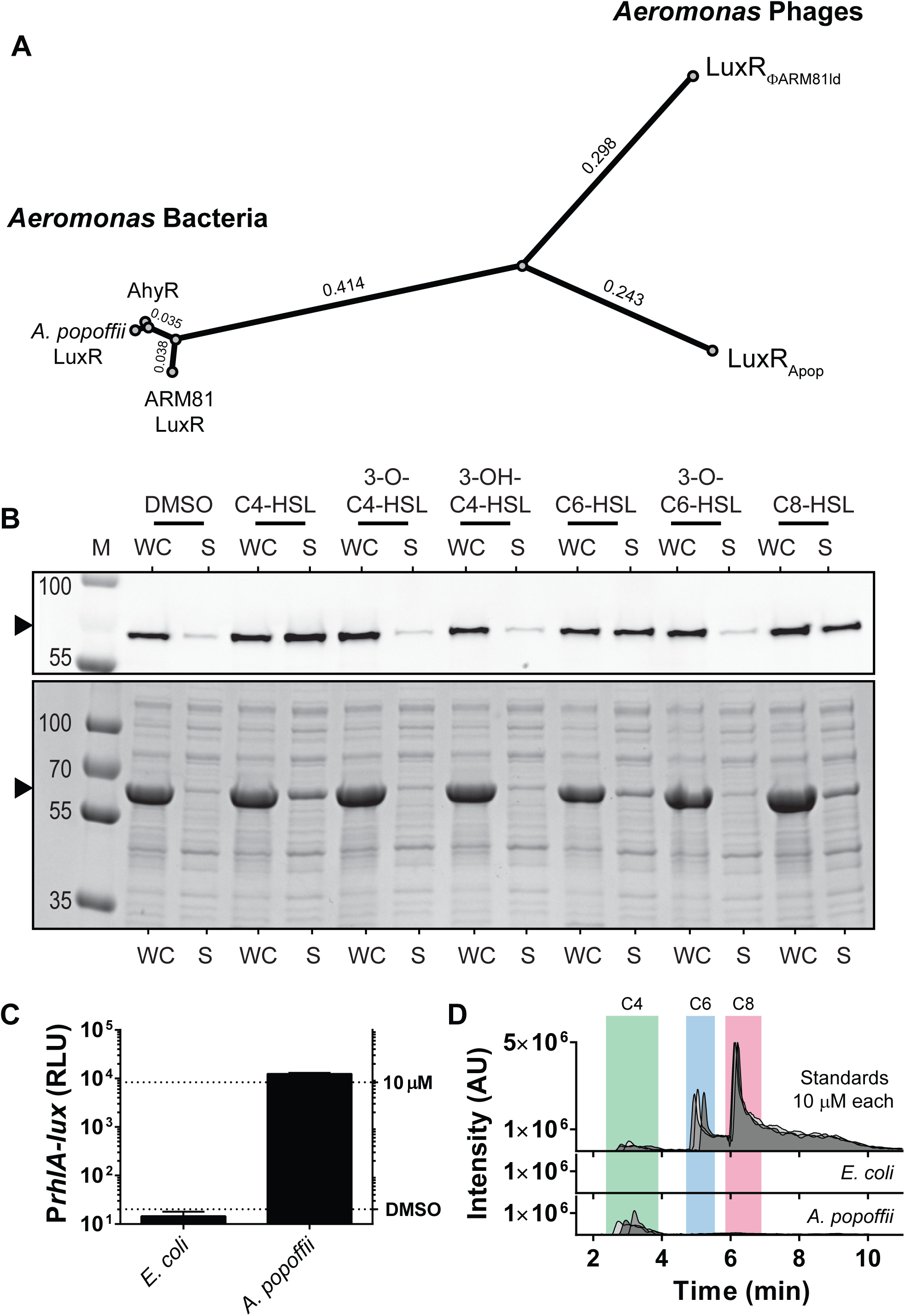
Host and phage harbor distinct LuxR proteins and the phage LuxR_Apop_ protein is solubilized by the AI produced by its host. (A) Phylogenetic analysis of three bacterially-encoded LuxR proteins (AhyR from *A. hydrophila* and the LuxR-type receptors from *Aeromonas sp*. ARM81 and *A. popoffii*) and the phage-encoded LuxR-type receptors (LuxR_ΦARM81ld_ and LuxR_Apop_). Branch lengths are indicated. (B) Western blot (upper) and total protein (lower) showing HALO-LuxR_Apop_ in the whole cell (WC) lysate and soluble (S) fractions from recombinant *E. coli* supplied with 75 µM of the indicated AHL or an equivalent volume of DMSO. Arrowheads indicate HALO-LuxR_ΦApop_. M denotes marker (PageRuler Plus; representative bands are labeled). (C) Bioassay to detect C4-HSL based on the *Pseudomonas aeruginosa* RhlR receptor and the target *rhlA* promoter. The *E. coli* bioassay strain does not produce C4-HSL. Thus, P*rhlA-lux* expression depends on exogenous C4-HSL. Shown is P*rhlA-lux* activity following addition of 25% v/v of cell free culture fluids prepared from the indicated strains. Data are represented as mean ± std with *n*=3 biological replicates. The upper and lower dashed lines, respectively, designate the reporter activity in response to 10 µM C4-HSL and following addition of an equivalent volume of DMSO solvent. (D) LC-MS chromatogram of 10 µM C4-(green), C6-(blue), and C8-HSL (red) (172.097, 200.128, 228.159 m/z, respectively) standards in filtered LB medium and the same analyses of cell free culture fluids prepared from*E. coli* and *A. popoffii*. The data show 3 biological replicates for culture fluids and 3 technical replicates for standards.

We do not yet know what target promoters are controlled by these phage LuxR receptors nor how QS drives phage biology. In the characterized vibriophage-*Vibrio* case, addition of AI causes lysis when it is bound by the phage QS receptor. We found that addition of C4-, C6-, or C8-HSL to *A. popoffii* does not induce lysis (Figure S3), and, as mentioned, we were unable to obtain the *Aeromonas sp*. ARM81 host strain to test for a similar effect with phage ΦARM81ld. We suspect that, beyond the AI, the phage ΦARM81ld and Apop QS receptor controlled pathways could require some additional input that is not present in our experiments. Alternatively, the phage QS receptor controlled pathways could regulate a different aspect of phage-host or phage-phage biology, for example, superinfection exclusion. We are testing such possibilities now.

The Apop LuxR-type receptor was revealed through analysis of sequencing data of its host, *A. popoffii*, a bacterium not previously known to harbor a phage. We take this finding as preliminary evidence that additional QS components are encoded on phage genomes and await discovery. It appears that, while different members of the same bacterial genus possess highly similar QS receptors, the QS receptors on their resident phages are more varied. Whether this finding hints at general principles underlying evolution of phage-encoded QS receptors can be resolved with the identification of additional examples. Our hypothesis is that the LuxR-type receptors allow the phages to eavesdrop on host bacterial communication and, using the information they glean, the phages optimally alter their biology. This hypothesis mirrors the mechanism described for the vibriophage-*Vibrio* case described above (4). Other phage-bacterial QS interactions are also possible. For example, the uncharacterized *Myoviridae sp*. mentioned above that was identified in our initial search for viral QS receptors, encodes a predicted AHL binding receptor, and located immediately upstream is an ORF predicted to encode a LuxI-type AI synthase (Figure S1A). This arrangement suggests a model similar to the phage-phage arbitrium system (3), but with the added complexity that the host could produce and/or respond to the same QS AI as the phage. These occurrences, primarily garnered from database searches, exemplify, as first substantiated by Hargreaves et al., that phage-bacterial QS interactions may be common (2). Future work could be aimed at identifying the regulons controlled by these phage QS components, whether they are phage-specific, host-specific, or whether they drive both phage and host biology, and their consequences to both the phage and the host.

## Methods

### Bacterial strains, plasmids, and growth conditions

Strains and plasmids used in this study are listed in Tables S1 and S2, respectively. *E. coli* strains and *A. popoffii* CIP 105493 were grown with aeration in Luria-Bertani (LB-Miller, BD-Difco) broth. Unless otherwise indicated, *E. coli* was grown at 37°C and *A. popoffii* at 30°C. Where appropriate, antibiotics were used at: 100 μg mL^−1^ ampicillin (Amp, Sigma), 100 μg mL^−1^ kanamycin (Kan, GoldBio), and 10 μg mL^−1^ chloramphenicol (Cm, Sigma). Inducers were used at: 200 ng mL^−1^ mitomycin C (MMC, Sigma), 0.1% L-arabinose (ara, Sigma), and 200 µM isopropyl β-D-1-thiogalactopyranoside (IPTG, GoldBio). Unless otherwise indicated, all autoinducers (AIs) were used at 75 μM. WuXi AppTec provided AIs except for 3-OH-C4-HSL which was made in house and C6-HSL which was obtained from Cayman Chemical.

### Cloning techniques

Primers used for plasmid construction and for generating the *luxR*_*ΦARM81ld*_ dsDNA gene block sequence are listed in Table S3 (Integrated DNA Technologies). The publicly available annotation of *luxR*_*ΦARM81ld*_ (*p37*) associated with ΦARM81ld (accession: KT898133) was used as the reference with the exception that, based on our own sequence alignments, the *luxR*_*ΦARM81ld*_ start codon is more likely 4 codons downstream from the annotated start site.

The T7-based vector for production of the two phage LuxR proteins was constructed so that both proteins had identical C-terminal linkers to a TEV cleavage site and HALO-HIS_6_ tag. The backbone for the expression vectors was prepared by PCR amplification of pJES-178, a vector derived from pH6HTC (Promega Corp [CAT#G8031]), with primers JSO-1268 x JSO-1435. The amplification strategy generated an N-terminal PacI site and a blunt C-terminal TEV-HALO-HIS_6_ tag. The amplified backbone product was treated with DpnI, CIP, and PacI. The *luxR*_*ΦARM81ld*_ insert was prepared by PCR amplification of JSgblock-93 with JSO-1438 x JSO-1440, followed by treatment with PacI and ligation into the backbone to generate pJES-179. The *luxR*_*Apop*_ insert was prepared by PCR amplification of total DNA extracted from *A. popoffii* CIP 105493 with JSO-1502 x JSO-1503 followed by treatment with PacI and ligation into the backbone to generate pJES-181. The Apop mini-plasmid, pJES-180, used to demonstrate the presence of the functional *repA* gene on the phage contig, was constructed by PCR amplification of total DNA extracted from *A. popoffii* with JSO-1452 x JSO-1462 followed by blunt-end ligation with a pRE112-derived PCR fragment (the product of amplification with JSO-931 x JSO-932) encoding Cm^R^ and the *pir*-dependent *oriR6ky* (12). The ligated product was transformed into *E. coli* TOP10 (Invitrogen), which lacks *pir*, so plasmid replication could only be accomplished via the cloned *repA* gene. The cloned *repA*-containing fragment could be amplified with primers JSO-1522 x JSO-1452.

Q5 High Fidelity Polymerase, T4 DNA ligase, and restriction enzymes were obtained from NEB. Constructs were transformed into TOP10 *E. coli*, where they were isolated and verified by sequencing. All DNA was introduced by electroporation using 0.1 cm gap cuvettes (USA Scientific) with a Bio-Rad MicroPulser.

### Total protein and HALO Western blot to assess phage LuxR solubility

The pJES-179 and pJES-181 plasmids carrying HALO-LuxR_ΦARM81ld_ and HALO-LuxR_Apop_, respectively, were transformed into *E. coli* T7Express *lysY/I*^*q*^ (NEB). Overnight cultures were back-diluted 1:100 into 100 mL fresh medium and grown at 34°C to OD_600_ ∼ 0.4-0.6. IPTG was added to the cultures and subsequently, they were divided into seven 10 mL aliquots. Six of the aliquots received 75 μM of an AI, and the seventh aliquot received an equivalent volume (0.1% v/v) of DMSO. The cultures were shaken at 34°C for an additional 8 h. The cells were pelleted at 4,000 RPM x 10 min and stored at −80°C until further use.

For protein solubility assessment, each pellet was thawed, resuspended and lysed in 200 μL of HALO-BugBuster lysis buffer (1x BugBuster (Millipore) supplemented with 100 mM NaCl, 1 mM DTT, 1x Halt protease inhibitor cocktail (Thermo), 0.5 mM EDTA, 3.5 μM HaloTag-Alexa Fluor 660 ligand (HALO-Alexa_660_, Promega), and 2 μL mL^−1^ lysonase bioprocessing reagent (Millipore)). After 30 min incubation at 30°C, half of each lysate was transferred to a new tube on ice (designated Whole Cell lysate fraction, WC) and the remainder was subjected to centrifugation at 15,000 g for 30 min at 4°C and the clarified supernatant recovered (designated Soluble fraction, S). 2 μL of each WC sample was diluted into 28 μL of lysis buffer (15-fold dilution), and 5 μL of each S sample was diluted into 25 μL of lysis buffer (6-fold dilution). 10 μL of 4x Laemmli sample buffer (Bio-Rad) was added to each tube, followed by incubation at 70°C for 15-20 min. 10 μL of each sample was separated by SDS-PAGE in 4–20% Mini-Protein TGX gels (Bio-Rad) and stained with Coomassie dye for total protein. A second gel, run in parallel, was loaded with the above WC and S samples diluted a further 15-fold and 6-fold, respectively. HALO-Alexa_660_ was imaged using the Cy5 setting of an ImageQuant LAS 4000 (GE). Exposure times never exceeded 15 sec. PageRuler Plus prestained protein ladder (Thermo) was used as the marker.

### *A. popoffii* growth and lysis assays

Growth: Four overnight cultures of *A. popoffii* were back-diluted 1:1,000 into fresh LB broth and quadruplicate 198 µL aliquots from each culture were dispensed into wells of a 96-well plate containing 2 µL of 10 mM C4-, C6-, C8-HSL or an equivalent volume of DMSO. Lysis: Four overnight cultures of *A. popoffii*, each prepared from a different single colony, were back-diluted 1:100 into LB broth, grown to OD_600_ = 1.0, and again back-diluted to OD_600_ = 0.15. From each culture, duplicate aliquots of 198 µL were dispensed into wells of a 96-well plate (Corning Costar, CAT#3904) containing 2 µL MMC or water. In both the growth and lysis assays, plates were shaken at 30°C and a BioTek Synergy Neo2 Multi-Mode reader was used to measure OD_600_ every 7.5 min. For the DNase assay, 1 mL of *A. popoffii* culture grown and induced with MMC as described above was allowed to lyse for 5 h at 30°C. The sample was subjected to centrifugation at 15,000 g for 2 min, the supernatant recovered, and DNase (DNA-free Kit, Thermo) added. After incubation at 37°C for 40 min followed by DNase inactivation, DNA was purified using the Phage DNA isolation kit (Norgen Biotek). PCR amplification of Apop DNA was carried out using primers JSO-1514 x JSO-1503, and the product verified by electrophoresis and sequence analysis.

### Bioassay for *A. popoffii* produced AHLs

The bioassay to measure C4-HSL in cell-free culture fluids employed a previously published *E. coli* reporter strain (JP-117) harboring a plasmid with P*rhlA-lux* and a plasmid with pBAD-*rhlR* (13). An overnight culture of JP-117, grown at 30°C, was back-diluted 1:100 in LB broth containing 0.1% L-arabinose and returned to growth at 30°C for 2 h prior to being dispensed into a 96 well plate (150 µL per well). Cell-free culture fluids from an overnight culture of *A. popoffii* or a non-AHL producing negative control (*E. coli* T7Express *lysY/I*^*q*^) were prepared by centrifugation (1 min × 15,000 RPM) followed by filtration through 0.22 μm Costar Spin-X tubes (Corning). 50 μL of the resulting cell-free culture fluids were added to the wells. For a positive control, a 40 μM synthetic C4-HSL standard was made in LB, and 50 µL of this preparation was added to 150 μL aliquots of the JP-117 reporter strain. Bioluminescence and OD_600_ were measured in a BioTek Synergy Neo2 Multi-Mode reader 7 h later. Relative light units were calculated by dividing the bioluminescence by the OD_600_ of the culture.

### Liquid chromatography-mass spectrometry to detect C4-, C6-, and C8-HSL

Cell-free fluids were prepared from overnight cultures of *A. popoffii* or *E. coli* T7Express *lysY/I*^*q*^ (negative control) as described above. Standards containing synthetic C4-, C6- and, C8-HSL (each at 10 μM) were made in LB broth. Culture fluids and standards were loaded onto a 1 mm × 75 mm C18 column (ACE 3 C18 PFP, Mac-Mod) using a Shimadzu HPLC system and PAL auto-sampler (20 μL per injection) at a flow rate of 70 μL min^−1^, using a previously published method (4). Full scan MS data were acquired with a LTQ-Orbitrap XL mass spectrometer (Thermo) at a resolution of 30,000 in profile mode from the m/z range of 170 – 240. C4-, C6-, and C8-HSL were detected by performing XIC at m/z of 172.09736 (C_8_H_14_NO_3_), 200.128668 (C_10_H_18_NO_3_), and 228.15996 (C_12_H_22_NO_3_), respectively, each with a mass accuracy of ± 10 ppm. Caffeine (2 µM in 50% acetonitrile with 0.1% formic acid) was injected as a lock mass using an HPLC pump (LC Packing) with a flow splitter and delivered at an effective flow rate of 20 µL min^−1^ through a tee at the column outlet. Files were processed in Xcalibur (Thermo).

### Quantification and statistical analysis

Data were recorded, analyzed, and plotted using Microsoft Excel and GraphPad Prism 6. Sequencing results, alignments, phylogenetic trees, plasmid maps, and genomes were assembled using SnapGene (GSL Biotech), UGENE (Unipro), Geneious (Biomatters Limited), and ApE (M. Wayne Davis). Whole genome alignment was performed using Mauve (14). The phage-like contig from *A. popoffii* CIP 105493 and the contig containing the *Aeromonas* host *luxR* locus are available at NCBI (accession: NZ_CDBI01000063.1 and NZ_CDBI01000091.1, respectively). The genome sequence of the uncharacterized Myovirdiae phage (Figure S1A) is available at NCBI (accession: MH622937.1) as “Myoviridae sp. isolate ctcg_2”. PHASTER analysis was performed at http://phaster.ca/ using the *A. popoffii* contig as the input (10).

### Data and software availability

All experimental data that support the findings of this study are available from the corresponding author upon request.

## Acknowledgments

This work was supported by the Howard Hughes Medical Institute, NIH Grant 2R37GM065859, National Science Foundation Grant MCB-1713731 and the Max Planck Society-Alexander von Humboldt Foundation (to B.L.B.). J.E.S. was supported by the Department of Defense (DoD) through the National Defense Science & Engineering Graduate Fellowship (NDSEG) Program. We thank Saw Kyin for work in the Princeton Proteomics & Mass Spectrometry Core, Christopher Buck for insight on the *Myovirdiae* sp. phage, and all members of the Bassler lab for thoughtful discussions.

## Author Contributions

J.E.S. and B.L.B. conceptualized the project. J.E.S. performed all experiments. J.E.S. and B.L.B. analyzed the data; and J.E.S. and B.L.B. wrote the paper.

## Declaration of Interests

The authors declare no competing financial interests.

## References

1. Papenfort K, Bassler BL. 2016. Quorum sensing signal–response systems in Gram-negative bacteria. Nat Rev Microbiol 14:576–588.

2. Hargreaves KR, Kropinski AM, Clokie MRJ. 2014. What Does the Talking?: Quorum Sensing Signalling Genes Discovered in a Bacteriophage Genome. PLoS ONE 9.

3. Erez Z, Steinberger-Levy I, Shamir M, Doron S, Stokar-Avihail A, Peleg Y, Melamed S, Leavitt A, Savidor A, Albeck S, Amitai G, Sorek R. 2017. Communication between viruses guides lysis–lysogeny decisions. Nature 541:488–493.

4. Silpe JE, Bassler BL. 2019. A Host-Produced Quorum-Sensing Autoinducer Controls a Phage Lysis-Lysogeny Decision. Cell 176:268–280.e13.

5. Papenfort K, Silpe JE, Schramma KR, Cong J-P, Seyedsayamdost MR, Bassler BL. 2017. A Vibrio cholerae autoinducer–receptor pair that controls biofilm formation. Nat Chem Biol 13:551–557.

6. Dziewit L, Radlinska M. 2016. Two novel temperate bacteriophages co-existing in Aeromonas sp. ARM81 – characterization of their genomes, proteomes and DNA methyltransferases. J Gen Virol 97:2008–2022.

7. Swem LR, Swem DL, O’Loughlin CT, Gatmaitan R, Zhao B, Ulrich SM, Bassler BL. 2009. A Quorum-Sensing Antagonist Targets Both Membrane-Bound and Cytoplasmic Receptors and Controls Bacterial Pathogenicity. Mol Cell 35:143–153.

8. Swift S, Karlyshev AV, Fish L, Durant EL, Winson MK, Chhabra SR, Williams P, Macintyre S, Stewart GS. 1997. Quorum sensing in Aeromonas hydrophila and Aeromonas salmonicida: identification of the LuxRI homologs AhyRI and AsaRI and their cognate N-acylhomoserine lactone signal molecules. J Bacteriol 179:5271–5281.

9. Huys G, Kámpfer P, Altwegg M, Kersters I, Lamb A, Coopman R, Lüthy-Hottenstein J, Vancanneyt M, Janssen P, Kersters K. 1997. Aeromonas popoffii sp. nov., a Mesophilic Bacterium Isolated from Drinking Water Production Plants and Reservoirs. Int J Syst Evol Microbiol 47:1165–1171.

10. Arndt D, Grant JR, Marcu A, Sajed T, Pon A, Liang Y, Wishart DS. 2016. PHASTER: a better, faster version of the PHAST phage search tool. Nucleic Acids Res 44:W16–W2

11. Ortori CA, Dubern J-F, Chhabra SR, Cámara M, Hardie K, Williams P, Barrett DA. 2011. Simultaneous quantitative profiling of N-acyl-l-homoserine lactone and 2-alkyl-4(1H)-quinolone families of quorum-sensing signaling molecules using LC-MS/MS. Anal Bioanal Chem 399:839–850.

12. Edwards RA, Keller LH, Schifferli DM. 1998. Improved allelic exchange vectors and their use to analyze 987P fimbria gene expression. Gene 207:149–157.

13. Paczkowski JE, Mukherjee S, McCready AR, Cong J-P, Aquino CJ, Kim H, Henke BR, Smith CD, Bassler BL. 2017. Flavonoids Suppress *Pseudomonas aeruginosa* Virulence through Allosteric Inhibition of Quorum-sensing Receptors. J Biol Chem 292:4064–4076.

14. Darling ACE. 2004. Mauve: Multiple Alignment of Conserved Genomic Sequence With Rearrangements. Genome Res 14:1394–1403.

