## supplemental figures and table for "Phage-Encoded LuxR-Type Receptors Responsive to Host-Produced Bacterial Quorum-Sensing Autoinducers"

SILPE & BASSLER

### FIGURE S1

## A

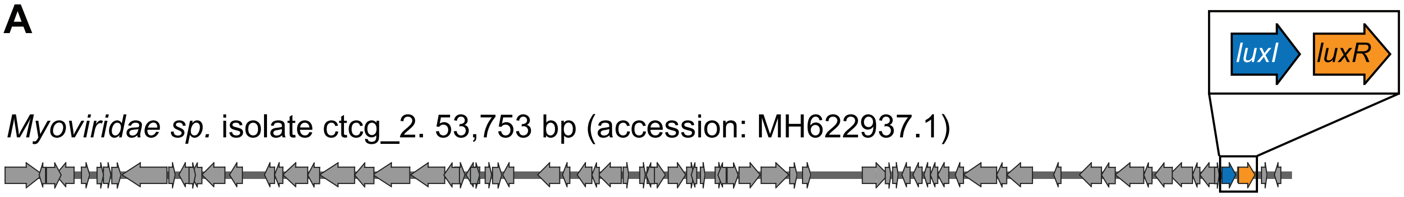


## B

```
                                 1        10        20        30
Consensus                        MKXXXJLEYLEHFTXVTDXDRLXELIGRFTLGMGYDXX
Aeromonas phage ΦARM81ld LuxR    MN---IEHHFSRFDAVTSEQGLTAEISAFAASLGIDQF
A. popoffii LuxR homolog 1       MKQDQLLEYLEHFTLVTDGDRLAELIGRFTLGMGYDYY
A. popoffii LuxR homolog 2       MD---IKRVIDKFSVVTTEGELTRALASFSSAMGFHQF
A. hydrophila AhyR               MKQDQLLEYLEHFTSVTDGDRLAELIGRFTLGMGYDYY

                                   40        50        60        70
Consensus                        RFALIIPMSXQRPKXVJFXXCXXSWVXAYTANHMLACD
Aeromonas phage ΦARM81ld LuxR    RFALLIPSSLAKPRAVIFSHCSEAWVAEYASAGLLRID
A. popoffii LuxR homolog 1       RFALIIPMSMQRPKVVLFNQCPDSWVQAYTANHMLACD
A. popoffii LuxR homolog 2       RLALIIPQSLQRPIAVIFSHCSEEWVAQYTKENMLQRD
A. hydrophila AhyR               RFALIIPMSMQRPKVVLFNQCPDSWVQAYTANHMLACD

                                     80        90        100       110
Consensus                        PIIXLARXQTLPIYWNRXXXRARFLXEGXXXVMXXAAE
Aeromonas phage ΦARM81ld LuxR    PIIHLALRQTRPIYWHSSLPHPRHLPPGAMEVMERAAS
A. popoffii LuxR homolog 1       PIIQLARKQTLPIYWNRLEGRARFLQEGSLDVMGLAAE
A. popoffii LuxR homolog 2       PIIHMAMRQTLPIFWQTSIIRTPDLPLGAMEVMERASE
A. hydrophila AhyR               PIIQLARKQTLPIYWNRLDERARFLQEGSLDVMGLAAE

                                       120       130       140       150
Consensus                        FGLRNGXSFPLXGAAGENGILSFITAXXXXSDLLLESS
Aeromonas phage ΦARM81ld LuxR    FGLRNGVSFPLRGARGEYGILSFVTKDIG-TAGLMEAS
A. popoffii LuxR homolog 1       FGLRNGISFPLHGAAGENGILSFITAEQASSDLLLDSS
A. popoffii LuxR homolog 2       YGLRNGVSFPLRGAGGEAGILSFITNDYG-SGHLLETS
A. hydrophila AhyR               FGLRNGISFPLHGAAGENGILSFITAERASSDLLLESS

                                         160       170       180       190
Consensus                        PILXXMXBYIFEAAIRXVXVXXXEDDPQEALTDRETEC
Aeromonas phage ΦARM81ld LuxR    PWLRLAADVIFESAIRVASF---GSPGNLALTRREKEC
A. popoffii LuxR homolog 1       PILSWMSNYIFEAAIRIVRVSLREDDPQEALTDRETEC
A. popoffii LuxR homolog 2       PLLRLVADYILDAAIRVVSK---RGQ-DRSLTKREVEC
A. hydrophila AhyR               PILSWMSNYIFEAAIRIVRVSLREDDPQEALTDRETEC

                                           200       210       220
Consensus                        LFWASEGKTSGEIAXILGITXRTVXYXJXQVTRKTGSM
Aeromonas phage ΦARM81ld LuxR    LAWASEGKTTTEIAAILGITPRTVTYYIQQVLGKTHST
A. popoffii LuxR homolog 1       LFWASEGKTSGEIACILGITERTVNYHLNQVTRKTGSM
A. popoffii LuxR homolog 2       LFWASEGKTAAEIGAILGIVPRTVTYYIQEAVTKTNSV
A. hydrophila AhyR               LFWASEGKTSGEIACILGITERTVNYHLNQVTRKTGSM

                                   230       240       250       260
Consensus                        NRXQAIAKXVSXGILLPNLEQVXVTNFPKLMQ
Aeromonas phage ΦARM81ld LuxR    NRDQAIAKAMAGGVLLPSLD-VDVDNVY
A. popoffii LuxR homolog 1       NRYQAIAKGVSSGILLPNLEQVLVTNFPKLMQ
A. popoffii LuxR homolog 2       NRDQAIAKAAMGGLLLPNLELVRVEDYL
A. hydrophila AhyR               NRYQAIAKGVSSGILLPNLEQVVVTNFPKLMQ
```

#### Supplemental figures

**Figure S1. The uncharacterized *Myoviridiae* phage encodes a LuxI-LuxR pair and *A. popoffii* encodes multiple LuxR-type QS receptors.**

(A) Genome organization of the uncharacterized *Myoviridiae* phage predicted to encode *luxI* and *luxR* genes (blue and orange, respectively). (B) Amino acid sequence alignment of $\text{LuxR}_{\Phi\text{ARM81ld}}$ with LuxR homologs from *A. popoffii* and the characterized AhyR QS receptor of *A. hydrophila*. *A. popoffii* homologs 1 and 2 represent the host LuxR receptor and the phage-encoded LuxR $\text{LuxR}_{\text{Apop}}$ receptor, respectively. Sites of high, low, and no identity are shaded black, gray, and white, respectively. Residues are numbered according to their positions in the consensus sequence.

SILPE & BASSLER

**FIGURE S2**

**A**

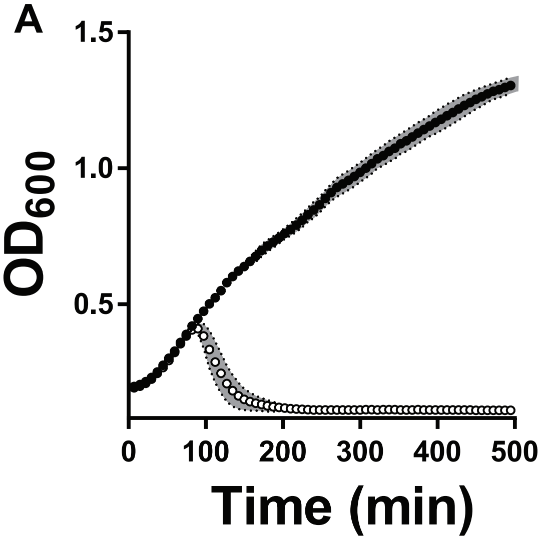


**B**

***Aeromonas* Bacterial LuxR Proteins (91% identity)**

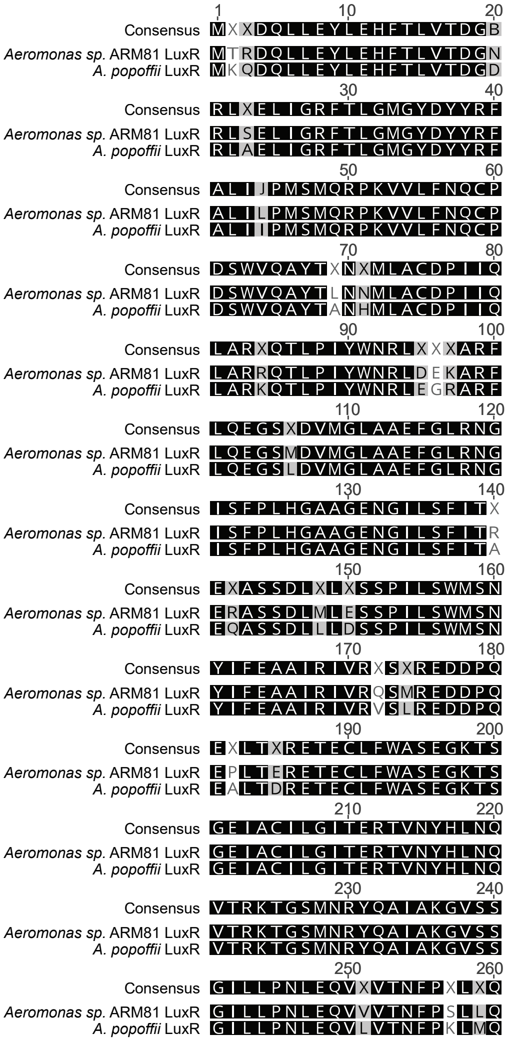


**C**

***Aeromonas* Phage LuxR Proteins (60% identity)**

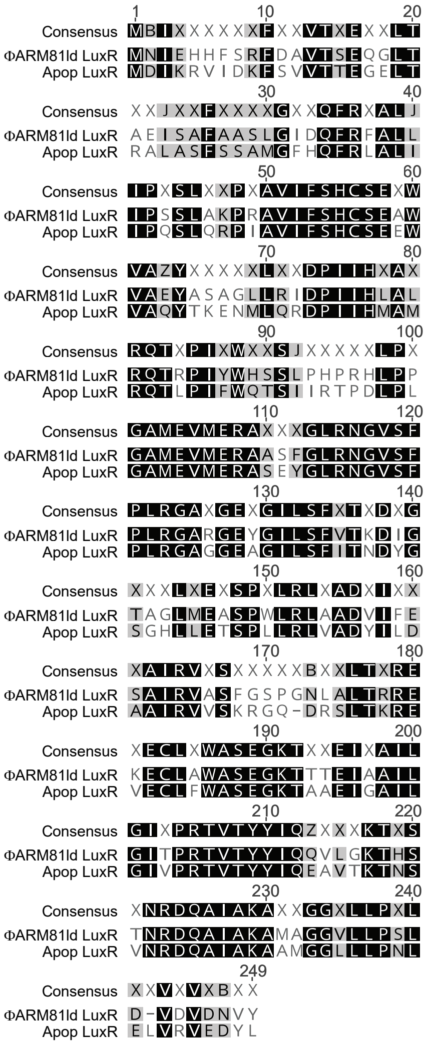

**Figure S2. MMC induces lysis of *A. popoffii* and host LuxR-type receptors are more similar to each other than to phage LuxR-type receptors.**

(A) Growth curve of *A. popoffii* strain CIP 105493 in the absence (black) or presence (white) of 200 ng $mL^{-1}$ MMC. (B) Amino acid sequence alignment of the *Aeromonas sp.* ARM81 and *A. popoffii* LuxR-type receptors. (C) Amino acid sequence alignment of the phage ΦARM81ld and Apop LuxR-type receptors. In (B) and (C), black and gray shading show identical and similar residues, respectively. Residues are numbered according to their positions in the consensus sequence.

**FIGURE S3**

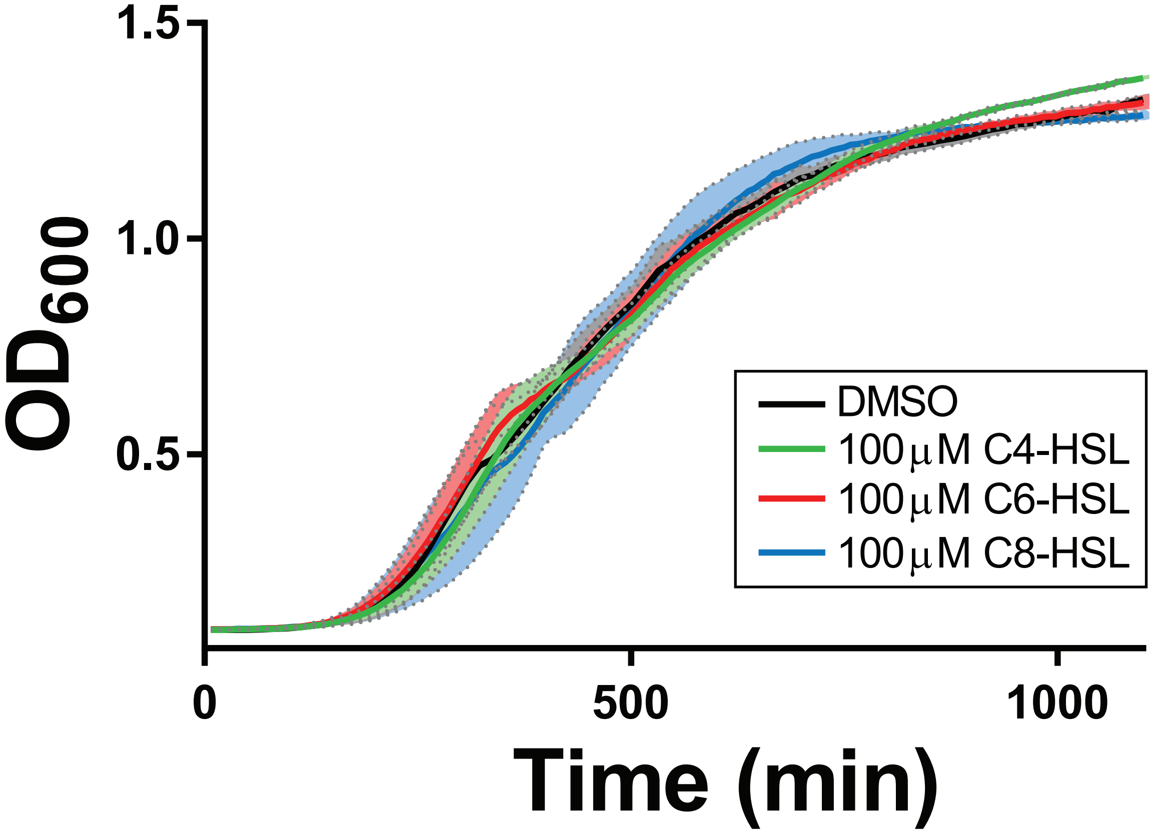
1.5
1.0
0.5
OD600
0
500
1000
Time (min)
DMSO
100 μM C4-HSL
100 μM C6-HSL
100 μM C8-HSL

**Figure S3. Exogenous addition of HSLs to *A. popoffii* does not induce lysis.** Growth curve of *A. popoffii* supplemented with the indicated HSL or an equivalent volume of DMSO. Data are represented as mean ± std with *n*=4 biological replicates.

#### Table S1: Bacterial strains used in this study

| Strain | Genotype | Reference |
|---|---|---|
| *E. coli* TOP10 | *F– mcrA Δ(mrr-hsdRMS-mcrBC) φ80lacZΔM15 ΔlacX74 recA1 araD139 Δ(ara-leu)7697 galU galK λ– rpsL(Str$^R$) endA1 nupG* | Invitrogen |
| *E. coli* T7Express *lysY/I$^q$* | *E. coli str. B, MiniF lysY lacIq(Cam$^R$) / fhuA2 lacZ::T7 gene1 [lon] ompT gal sulA11 R(mcr-73::miniTn10-Tet$^S$)2 [dcm] R(zgb-210::Tn10-Tet$^S$) endA1 Δ(mcrC-mrr) 114::IS10* | NEB |
| *A. popoffii* CIP 105493 | Wild-type, LMG 17541; BAA-243; CIP 105493 | ATCC BAA-243 |

#### Table S2: Plasmids used in this study

| # | Plasmid name (informal) | Plasmid ID | Strain ID (formal) | Relevant fragment | Marker, Origin | Source |
|---|---|---|---|---|---|---|
| 1 | pH6HTC-pT7-*HIS-HALO-cI*$_{VP882}$ | pJES-178 | JSS-1850 | *HIS-HALO-cI*$_{VP882}$ | Amp, pBR322 | This study |
| 2 | pH6HTC-pT7-*HIS-HALO-LuxR*$_{\Phi ARM81ld}$ | pJES-179 | JSS-1868 | *HIS-HALO-luxR*$_{\Phi ARM81ld}$ | Amp, pBR322 | This study |
| 3 | *repA*$_{Apop}$ | pJES-180 | JSS-1930 | *repA*$_{Apop}$ | Cm, Apop and oriR6ky | This study |
| 4 | pH6HTC-pT7-*HIS-HALO-LuxR*$_{Apop}$ | pJES-181 | JSS-1950 | *HIS-HALO-luxR*$_{Apop}$ | Amp, pBR322 | This study |
| 5 | pBAD-*RhlR* | pJP-2 | JP-117 / BB-0386 | *rhlR* | AmpR, pBR322 | (1) |
| 6 | P*rhlA-lux* | pJP-11 | JP-117 / BB-0386 | P*rhlA-lux* reporter | KanR, pSC101 | (1) |

##### Supplemental references for Table S2.


1. Paczkowski JE, Mukherjee S, McCready AR, Cong J-P, Aquino CJ, Kim H, Henke BR, Smith CD, Bassler BL. 2017. Flavonoids Suppress *Pseudomonas aeruginosa* Virulence through Allosteric Inhibition of Quorum-sensing Receptors. J Biol Chem 292:4064–4076.

#### Table S3: Oligonucleotides and dsDNA used in this study

| Primer | Sequence (5' - 3') | 5' Mod* |
|---|---|---|
| JSO-1268 | GAGCCAACCACTGAGGATCT | |
| JSO-1435 | GACGTTACCAAAATTCATCATTAATTAACCTCCTG | |
| JSO-1438 | GTTTTTTAATTAATTGGCCGATGATATGAACATAGAGCAC | |
| JSO-1440 | GTAGACGTTGTCGACATCCACG | P' |
| JSO-1502 | GTTTTTTAATTAATGGACATCAAACGGGTTATCGATAAATTCA | |
| JSO-1503 | CAGATAATCCTCTACTCTGACCAGCT | P' |
| JSO-1452 | CCTCGATCGTTTAATCCACTCGATAGA | |
| JSO-1462 | AGAGCTATCAGGCTCATACACCC | |
| JSO-0931 | CTGTCTCTTATACACATCTTCTAGAAGAAGCTTGGGATC | P' |
| JSO-0932 | CTGTCTCTTATACACATCTCTGTTGCATGGGCATAAAG | P' |
| JSO-1522 | AGCCGAGTCCGTTTACCGG | |
| JSO-1514 | GACATCAAACGGGTTATCGATAAATTCAGC | P' |

| dsDNA | Sequence (5' - 3') |
|---|---|
| JSgblock-93 | GCCAACCACTGAGGATCTGTACTTTCAGAGCGATAACGCGGCCGATGATATGAACATAGAGC<br>ACCACTTCTCCCGCTTTGACGCGGTAACATCAGAACAAGGGCTGACAGCAGAGATTTCAGCG<br>TTTGCAGCCAGCCTCGGGATTGACCAGTTCCGCTTCGCCTTGCTCATCCCCTCGTCACTGGC<br>AAAGCCTCGGGCCGTCATTTTCAGCCACTGCAGTGAGGCTTGGGTGGCCGAATATGCCAGC<br>GCTGGCTTGCTTCGAATCGACCCTATCATCCATCTAGCACTGCGTCAGACCCGCCCCATTTAT<br>TGGCACTCAAGCCTGCCTCATCCTCGGCACCTCCCCCCAGGGGCAATGGAGGTCATGGAGC<br>GGGCTGCTTCTTTCGGCCTGCGCAACGGGGTGTCTTTTCCGCTGAGGGGGGCTCGAGGGGA<br>GTATGGGATCCTGTCGTTCGTGACGAAGGACATCGGCACTGCGGGCTTGATGGAGGCCAGT<br>CCCTGGCTTCGGCTGGCGGCTGACGTGATTTTTGAGTCGGCCATTCGGGTGGCATCGTTCG<br>GAAGCCCCGGAAACCTGGCCCTGACCCGCCGCGAAAAGGAGTGCCTGGCGTGGGCCAGTG<br>AGGGCAAGACGACCACCGAGATTGCCGCTATTCTTGGCATCACCCCCAGGACGGTGACCTAT<br>TACATTCAGCAGGTACTGGGGAAGACCCACAGCACGAACCGGGATCAGGCGATTGCCAAGG<br>CGATGGCCGGCGGTGTGCTGCTTCCTAGCCTGGACGTGGATGTCGACAACGTCTACTGATTA<br>ATTAACCAATTCCTGCAGGATTTTGCGGCCGCTTGCT |

*indicates phosphorylation at the 5’ end
